# A bibliometric analysis of gender in microbiology collaborations

**DOI:** 10.1101/2022.12.14.520436

**Authors:** Rachel M. Wheatley, Lois Ogunlana

## Abstract

Women are underrepresented in senior academic positions within microbiology globally. Studies show that gender bias affects the progression of women in academia, but there is evidence that improving conscious awareness of bias can improve equity in this regard. Here we carry out a bibliometric analysis of review articles within the microbiology field to investigate the statistical associations with author gender. We analyse the publication data from 1857 review articles published between 2010 and 2022 in three leading microbiology review journals: Nature Reviews Microbiology, Trends in Microbiology, and Annual Review of Microbiology. We find a significant association between the gender of the lead author and the gender of co-authors in multi-author publications. Review articles with men lead authors have a significantly reduced proportion of women co-authors compared to reviews with women lead authors. Given the existing differences in the proportions of men and women in lead author positions, this association may have important consequences for the relative visibility of women in microbiology, along with potential negative impacts on scientific output relating to reduced collaboration diversity. We further probe associations between gender and citation metrics, acknowledgement of contributions, and publishing during the Covid-19 pandemic within microbiology reviews.

## Introduction

Senior microbiology research positions within academia often show a significant underrepresentation of women. The percentage of doctorates awarded to women in the life sciences is over 50%, but the number of women in postdoctoral and tenure-track positions is less than 40% and 30%, respectively (1, 2). Only 18% of full professors in biology-related fields are women (3). Numerous studies have shown how gender bias exists in professional evaluations (4, 5), promotions (6), grant proposal success (7, 8), salaries (9, 10), and the acknowledgement of contributions to work (11). Improving conscious awareness of where these inequalities exist is a first step towards improving equity in this regard (12). The quantitative analysis of data associated with academic publications, i.e. bibliometric analyses, is one way in which this can be achieved.

Bibliometric analyses often focus on primary research. These types of analysis have revealed differences in manuscript submission outcomes (2), citation metrics (13, 14), and the volume of self-citations (15) associated with author gender. Such analyses typically neglect review articles, which can be considered a metric of who is an expert in the field. Authoring review articles can increase the profile, visibility, and citations of a researcher.

Although single author reviews are not uncommon, reviews in microbiology typically have multiple authors. To this end, review articles provide the opportunity to work collaboratively on an intellectual project, and the possibility to transcend some of the logistical barriers imposed on laboratory-based research. Studies of gender or diversity within groups working on collaborative tasks show that mixed-gender or otherwise diverse teams can produce better outputs (16-18) and higher quality science (19). Multi-author reviews can also provide senior researchers with the opportunity to contribute to the career development of their juniors. For example, sharing expertise and forming collaborations with other scientists during the review writing process may expand networks and increase profiles of researchers in their field. This outlines the importance of review publications in academic communities, and the significance of multi-author collaborations. But what decides how these collaborations form? Researchers choose collaborators based on expertise, but also social factors such as existing working structures or personal relationships play a role (20, 21). Homophily is the principle that similarity breeds connection between individuals (22), and gender homophily is the principle that individuals assort non-randomly with respect to gender. Due to this social structuring, it is possible that gender plays a role in the assembly of review collaborators (22).

In this study we carry out a bibliometric analysis of review articles in microbiology published between 2010 and 2022 in three leading microbiology review journals (Nature Reviews Microbiology, Trends in Microbiology, and Annual Review of Microbiology) to investigate the statistical associations with authorship gender. We investigate whether gender may play a role in the formation of microbiology collaborations by investigating whether there is an association between the gender of the lead author and the gender of co-authors in multi-author reviews. Given the existing differences in the proportions of men and women in lead author positions, an association may have important consequences for the relative visibility and career progression of women in microbiology, along with potential impacts on scientific output relating to reduced collaboration diversity (16-19). Finally, we take three known associations with gender in academic publications reported in the literature (citation metrics (13, 23-25), acknowledgment of contributions (11), and publishing during the Covid-19 pandemic (26-30)), and test for these associations within our microbiology dataset.

## Results and discussion

Publication data was downloaded from Web of Science for review articles published between January 2010 and June 2022 in Nature Microbiology Reviews, Trends in Microbiology, and Annual Review of Microbiology. The gender of authors was inferred from first names using Gender API, a social media-informed classification algorithm. This approach infers gender based on first names and has been used extensively in work examining gender in authorship of academic articles (e.g. (2, 13, 31, 32)). This produced a dataset of 1857 review papers with inferences on author gender (Supplementary Table 1). This dataset had a total of 5680 authors and a median of 3 authors per paper. All references to author gender (i.e. woman/man) in this study utilize inferred gender. We recognize the limitations of this approach. Inferred gender is distinct from the gender(s) that an individual may identify as, is restricted to a binary mode of inference (i.e. is not inclusive of non-binary individuals), and left a category of unknown inferences who were excluded from the downstream analysis.

We find there are over six times as many review articles published with author lists that were inferred to consist of exclusively men compared to exclusively women (Figure 1A, Figure 1B). Review articles with author lists that consist of exclusively men accounted for 43% of papers in the total dataset, compared to only 7% for exclusively women, and 54% for mixed gender (Figure 1A). These numbers include single author reviews, which account for 199/1857 publications.

**Figure 1.**
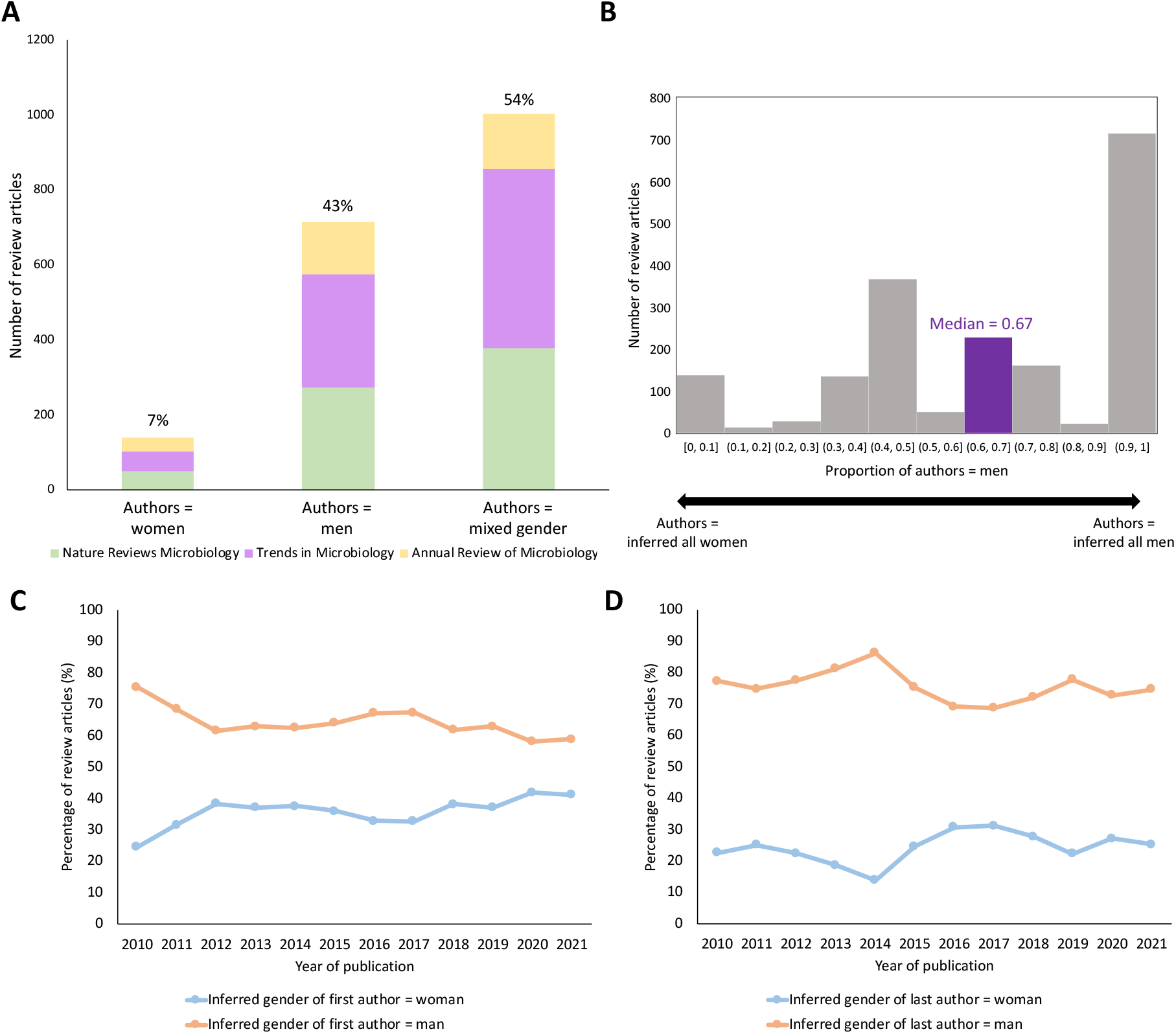
Overview of dataset. (A) Number of publications split by inferred gender of author list (authors = women, authors = men, authors = mixed gender) and journal (Nature Reviews Microbiology, Trends in Microbiology, Annual Review of Microbiology). Percentage rounded to 0 d.p. This number includes single author reviews, which account for 199/1857 publications. (B) Histogram showing the inferred gender of publication author-lists, as 0 = all authors inferred women and 1 = all authors inferred men. The median for this dataset is 0.67 (highlighted in purple). (C) Change over time in the proportion of men and women in first author positions. (D) Change over time in the proportion of men and women in last author positions.

Authorship positioning practices can be used to infer leadership roles within publications (33). In microbiology, as in other fields of biology, it is traditionally assumed that the first author has made the most contributions and the last author is the most senior scientist or principle investigator (34). Although there are some regional variations in the norms of authorship designation (35), by positioning conventions, the first or last named author should generally capture the majority of authors who have led or initiated a review article.

A total of 36% of reviews had women first authors. The proportion of publications with women first authors has increased over the last 10 years. Rising from 25% of reviews in 2010 to 41% of reviews in 2021 (Figure 1C), with the sharpest rise occurring between 2010 and 2012. The proportion of publications with women last authors has fluctuated, but from end points remained largely unchanged (Figure 1D). Women were last authors on 23% of reviews published in 2010 and 25% of reviews published in 2021 (Figure 1D). Overall a total of 24% of reviews had women last authors.

### Gender influences microbiology review collaborations

We wanted to investigate the relationship between the genders of the lead- and co-authors in multiauthor reviews (≥ 3 authors). Multi-author reviews (≥ 3 authors) accounted for 1066 publications in our dataset. Due to authorship positioning practices, the first or last named author should generally capture the majority of ‘lead authors’ (i.e. individuals most likely to have assembled publication contributions) for a review collaboration. To this end, we assessed the relationship between the inferred gender of first or last author with the inferred gender of co-authors on a publication.

We find a significant association between the inferred gender of the lead author and the inferred gender balance of co-authors in multi-author reviews (Figure 2, Table 1). On average 75% of coauthors were men when the first author was a man, compared to 67% of co-authors when the first author was a woman (Mann-Whitney, p < 0.01 two-tailed) (Figure 2A-B, Table 1). On average 67% of co-authors were men when the last author was a man, compared to 50% of co-authors when the last author was a woman (Mann-Whitney, p < 0.01 two-tailed) (Figure 2C-D, Table 1). This trend was observed across all three journals (Table 1).

**Figure 2.**
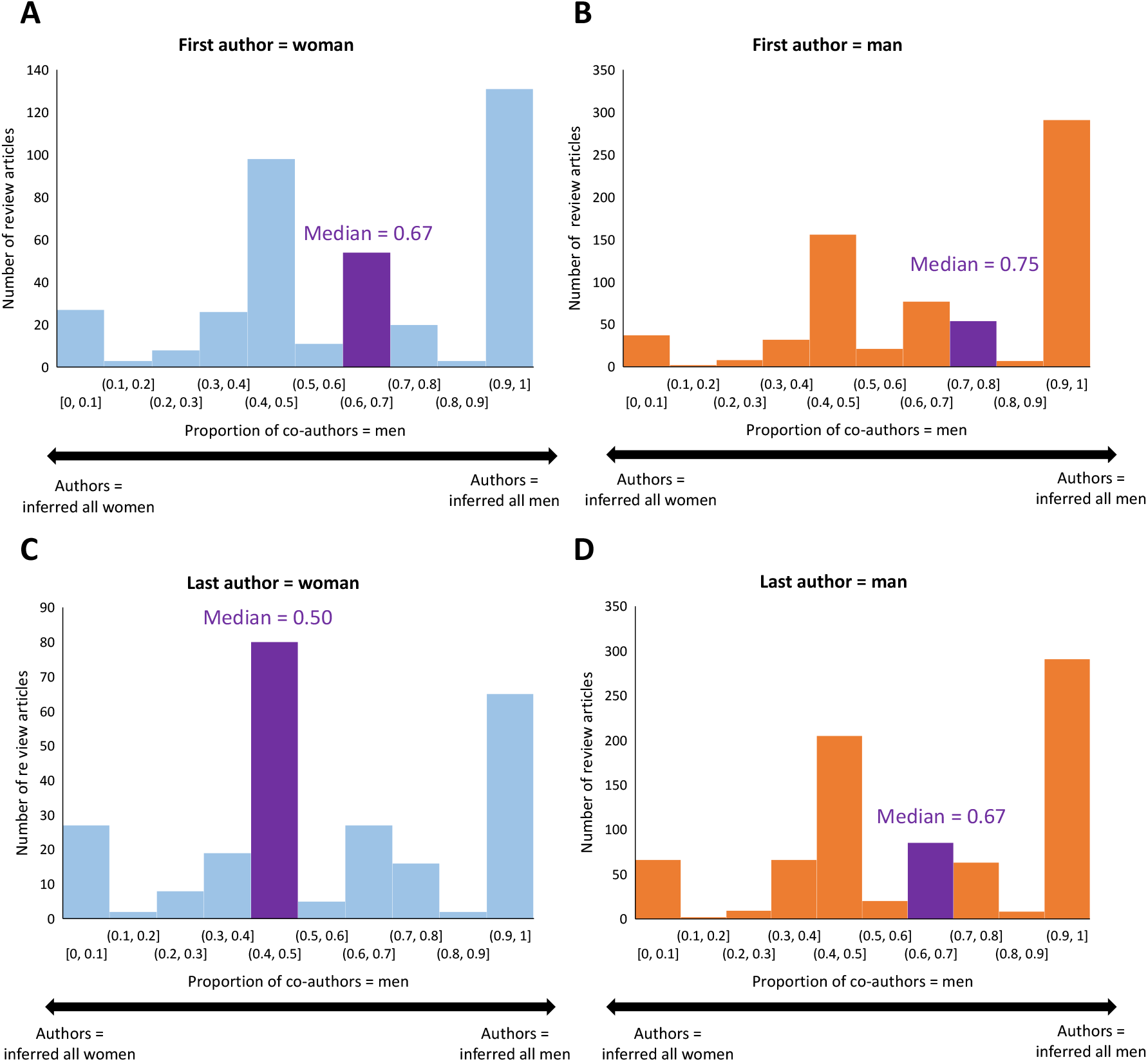
Gender distribution of co-authors by gender of first or last author. Shown for: (A) publications with women first authors (B) publications with men first authors (C) publications with women last authors (D) publications with men last authors. The bar containing the average (median) is highlighted in purple and median values are annotated on plots. Proportion of co-authors is shown on a scale where 1= all authors inferred men and 0= all authors inferred women.

**Table 1.** Inferred gender of first and last author and gender distribution of co-authors. On average 75% of co-authors were men when the first author was a man, compared to 67% of co-authors when the first author was a woman (Mann-Whitney, p < 0.01 two-tailed). On average 67% of co-authors were men when the last author was a man, compared to 50% of co-authors when the last author was a woman (Mann-Whitney, p < 0.01 two-tailed). For each journal and the total, the author row containing the highest proportion is highlighted in blue.

| Journal | Inferred gender of first author | Median proportion of co-authors = men | Total papers | Inferred gender of last author | Median proportion of co-authors = men | Total papers |
| --- | --- | --- | --- | --- | --- | --- |
| Annual Review of Microbiology | Woman | 0.50 | 52 | Woman | 0.50 | 38 |
|  | Man | 0.75 | 83 | Man | 0.67 | 97 |
| Nature Reviews Microbiology | Woman | 0.67 | 140 | Woman | 0.60 | 84 |
|  | Man | 0.75 | 271 | Man | 0.67 | 327 |
| Trends in Microbiology | Woman | 0.67 | 189 | Woman | 0.50 | 129 |
|  | Man | 0.67 | 331 | Man | 0.67 | 391 |
| Total | Woman | 0.67 | 381 | Woman | 0.50 | 251 |
|  | Man | 0.75 | 685 | Man | 0.67 | 815 |

We considered that as the number of authors on a publication increases, the probability of having mixed co-authorship should also increase. We confirmed that the size of the author list was not significantly different between publications with men or women lead authors (Mann-Whitney, p < 0.05 two-tailed). As a further consideration, we note a previous hypothesis has suggested that gender homophily in collaborations could be explained by subdivisions within scientific disciplines (36). They state that collaboration is based on disciplines or sub-communities within disciplines, which by chance can differ in gender (36). This would mean that collaborating with other researchers with similar expertise would result in an observed gender homophily in co-authorships when viewed across communities. Although we have not been able to test this in our dataset, by focusing on microbiology we are already focusing on a specific community within biology. Furthermore, testing of this hypothesis using large-scale datasets has reported that even when controlling for the differential structure of scholarly communities, significant associations between gender in co-authorship are found (36).

Overall our findings suggest that lead authors who are men are more likely to invite men co-authors to collaborate on review publications (Figure 2, Table 1). This association suggests gender is a factor that influences review collaboration formation in microbiology. This agrees with a number of studies across disciplines that shows collaborators assort non-randomly with respect to gender (35, 37-42), and supports that this gender homophily may be pervasive in microbiology publications.

In terms of impact, review articles can be considered a metric of who is an expert in the field, and authoring review articles can increase the profile, visibility and citations of a researcher. As such, given the overall differences in the proportions of men and women in lead author positions, this association may have important consequences for the relative visibility, profile and citations of women in microbiology. Furthermore, studies of gender or diversity within groups working on collaborative tasks show that mixed-gender or otherwise diverse teams can produce better outputs (16-18) and higher quality science (19). This suggests that reducing homophily within review collaborations could also be beneficial for scientific output (16-19).

### Association between author gender and citation metrics, acknowledgement of contributions, and publishing during the Covid-19 pandemic

Bibliometric analyses across several fields have reported that articles authored by women receive fewer citations than those written by men (13, 23-25). To investigate this within our microbiology dataset, we calculated the median number of citations for review articles by publication year and compared this to the inferred gender of first and last author (Figure 3A-B). For first author, we found that the median number of citations was higher for publications with men first authors for nine out of the twelve years compared (Figure 3A). However, this difference was only statistically significant for two of the years (Mann-Whitney, p < 0.05 two-tailed). For last author, we found that the median number of citations was higher for publications with men last authors for only seven out of the total twelve years compared (Figure 3B), and no differences in citations were statistically significant. While this overview shows limited clear significant differences, we recognise the limitations of this analysis. Our dataset is smaller compared to key citation studies across fields (13, 23-25), and limited to three high-impact journals. We additionally compared the top 20 most highly cited papers in our dataset. Out of these, only three (15%) had women last authors, which is lower than the overall proportion of papers with women last authors (24%) (Figure 3C).

**Figure 3.**
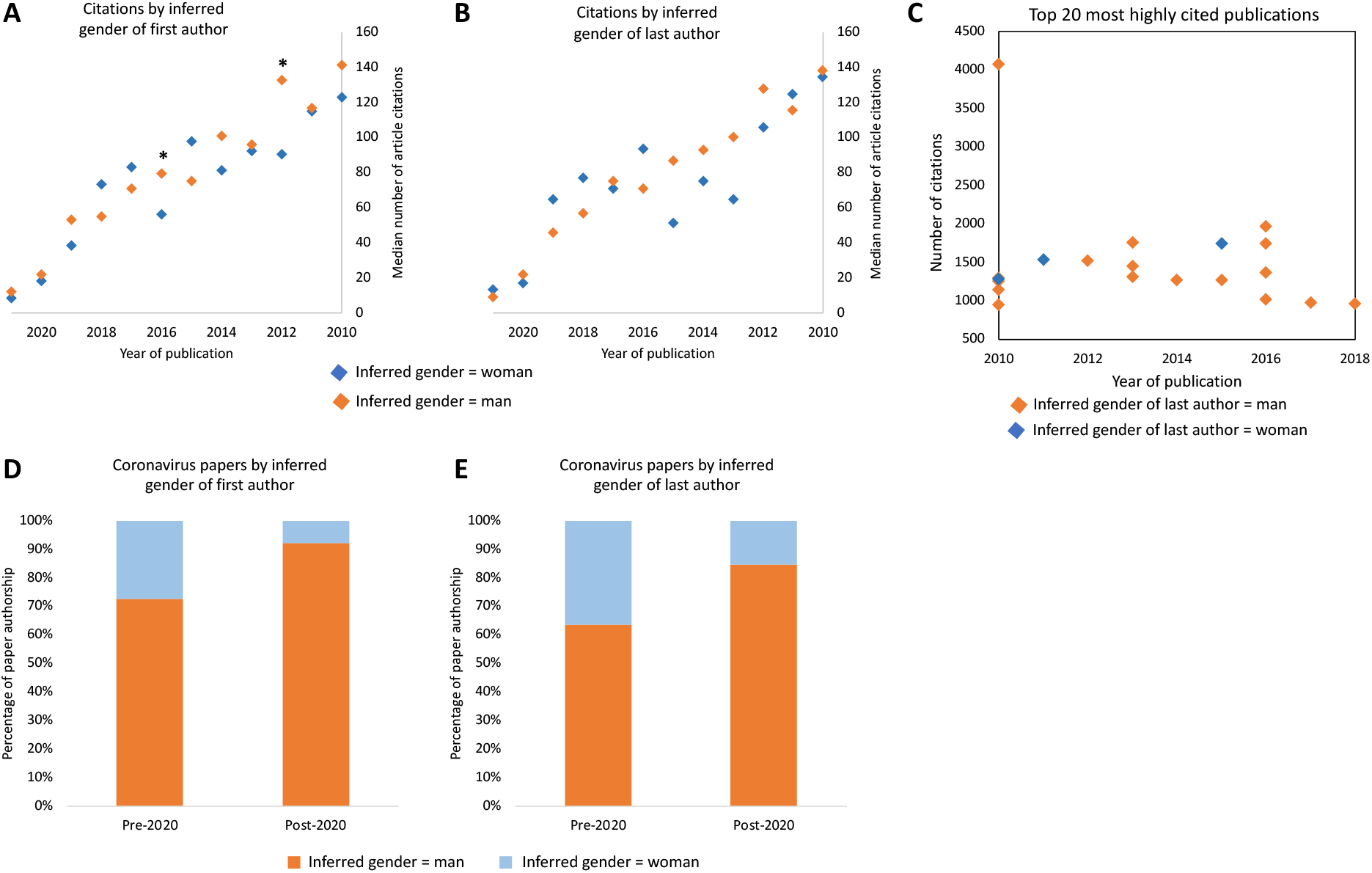
Citation metrics and coronavirus papers pre- and post-2020. (A) Median number of citations by publication year compared between papers by inferred gender of first author. Groups were compared by year, with differences significant in two years (* = Mann-Whitney, p < 0.05 two-tailed). (B) Median number of citations by publication year compared between papers by inferred gender of last author. Groups were compared by year and showed no statistically significant differences (MannWhitney, p < 0.05 two-tailed). (C) Number of citations for the top 20 most highly cited publications. (D) Coronavirus papers by inferred gender of first author pre-2020 and post-2020. (E) Coronavirus papers by inferred gender of last author pre-2020 and post-2020.

A number of studies have highlighted the disproportionate impact of the Covid-19 pandemic on women in academia (26-30). Surveys of academics in 2020 found that the strongest predictors of time lost from research were being female and the presence of young dependents (27). Analysis of primary literature has shown that the submission of papers by women in medical research decreased in 2020. Whereas submissions by men in the same field increased during this same period (29, 30). We wanted to investigate whether there was a relationship between timeline of the Covid-19 pandemic and publications in our dataset. We searched article keywords for instances of ‘coronavirus’. This identified pre-2020 reviews on non-SARS-COV-2 coronaviruses and post-2020 reviews on SARS-COV-2 coronaviruses. We found that the proportion of coronavirus reviews published by women decreased from pre-2020 to post-2020 (Figure 3D-E). From 27% of reviews pre-2020 to 8% of reviews post-2020 with women first authors, and from 36% of reviews pre-2020 to 15% of reviews post-2020 with women last authors (Figure 3D-E). Given we know that the primary research outputs on coronaviruses by women also decreased in this period (30), this finding is supportive of the disproportionate impact of the Covid-19 pandemic on women in microbiology as measured by publication outputs in the form of reviews during this period. We present this work with the caveat that microbiology review journals may not be the primary recipients for work relating to coronaviruses or the Covid-19 pandemic, and so this analysis was done on a small sample size of papers (26 papers) that limits the inferences that can be drawn.

Finally, a recent large-scale analysis of administrative data, surveys and interviews found that women in research teams are less likely to be credited for their contributions with authorship than their male peers (11). This is significant because any observed differences in scientific output relating to gender may in part be related to differences in attribution rather than differences in scientific contribution. We decided to explore this in our dataset by analysing the acknowledgements section of review papers and comparing this to author lists. As, there are multiple acknowledgement types, we focused on the contribution of figures or artwork because it is a quantifiable contribution that is likely more comparable between papers.

Within our dataset, a total of 59 review articles provided acknowledgement thanks for named individuals for figure or artwork contribution. Inferences on individual gender were successfully made in 55 of these cases. We find that roughly equal proportions of men and women (median = 0.50) were acknowledged in these acknowledgement sections. Author lists, on the other hand, are skewed towards higher proportions of men (median = 0.67). This difference was significant across the whole dataset (Mann-Whitney, p < 0.05 two-tailed) and directly within this sub-section of 55 review papers (Wilcoxon signed rank, p < 0.05 two-tailed). Although this type of analysis would benefit from more detailed data on author contributions, this seems to be in agreement with the findings of (11). However, we note that this result could also be influenced by figures being made by graduate researchers – a career stage where in biology genders are typically roughly equally distributed.

## Conclusions

In this study we carry out a bibliometric analysis of review articles in microbiology published between 2010 and 2022 in three leading microbiology review journals to investigate the statistical associations with authorship gender. Review articles can be considered a metric of experts in the field and can increase the profile and visibility of a researcher. They also provide the opportunity to work collaboratively on an intellectual project, and the possibility to transcend some of the logistical barriers imposed on laboratory-based research. The key finding of this work was that multi-author reviews with men as lead authors have a significantly reduced proportion of women co-authors compared to reviews with women as lead authors. Given the overall differences in the proportions of men and women in lead author positions, this may have important consequences for the relative visibility and progression of women in microbiology. Furthermore, this homophily may have negative impacts on the outputs produced in collaborations (16-19). We find women have an increased representation in acknowledgement sections compared to in authorship lists. Overview, our findings point to an underrepresentation of women in microbiology reviews, as has previously been shown to be the case in the publication of primary research (2). This study is an initial pilot study into the associations with authorship gender in microbiology publications, and supports the need for journals, editors, and researchers to consider the processes underlying the invitation or proposal of a review and assembly of collaborations, along with further research into this area.

## Methods

### Data collection and gender inference

Full article information was downloaded from Web of Science for all review articles published between January 2010 and June 2022 in Nature Reviews Microbiology, Trends in Microbiology, and Annual Review of Microbiology, with between one and ten authors. These three journals were chosen through a combination of impact factor evaluation and consensus-based discussion about the longterm professional impact of publication in these journals. This produced a dataset of 2025 papers. The gender of authors was inferred from first names using Gender API (43), a classification algorithm that uses self-reported gender from social media data. The data returned from Gender API included name, inferred gender (as “male”, “female”, or “unknown”), the number of instances the gender associated with the name was reported within the social media database, and a confidence of the gender inference (ranging from 0.5 to 1.0) based on this number of instances and the variability within them. Low accuracy assignments (below 0.7), unknowns, and instances where author first name was not provided in the author list were removed from the dataset. This produced a dataset of 1857 review papers with inferences on authorship gender (here referred to as woman/man) (Supplementary Table 1). Although this approach to infer gender from first name is used extensively in work examining gender in authorship of academic articles (e.g. (2, 13, 31, 32)), we recognize its limitations. Inferred gender based on first names is distinct from the gender(s) that an individual may self-identify as, is restricted to a binary mode of inference, and leaves a category of unknown gendered individuals who were downstream excluded from analysis. We are aware that names from certain countries, e.g. western European countries, are overrepresented in the Gender API database. This could mean nonwestern names may have a higher likelihood of being classified as unknown and therefore excluded from this analysis.

### Dataset analysis

Author inferred genders were converted into numerical (1=man, 0=woman), and this was used to calculate the gender distribution of co-authors within the author list of a paper. Publications with three or more authors were used to investigate the relationship between the inferred gender of the lead author and the inferred gender of co-authors in multi-author reviews. Data was tested for distribution normality using a Kolmogorov-Smirnov test. A non-parametric test was used to compare group distributions (Mann-Whitney, two-tailed), as done in (42). For analysis of citation metrics, the number of citations for reviews by publication year was compared to the inferred gender of first and last author (Mann-Whitney, two-tailed). To investigate the relationship between timeline of the Covid-19 pandemic and review publications we searched article keywords for instances of ‘coronavirus’. We used the results of this keyword search to compare number of pre-2020 reviews on non-SARS-COV-2 coronaviruses and number of post-2020 reviews on SARS-COV-2 coronaviruses by author gender. We systematically searched article acknowledgement sections for individuals attributed with figure or artwork contributions whose first name was given. As before, Gender API was used to infer the gender of individuals listed in these acknowledgements. The inferred gender of individuals listed in the acknowledgements was compared to the inferred gender of individuals listed in the author-list across the full dataset (Mann-Whitney, two-tailed) and within the sub-set of articles with included acknowledgements (Wilcoxon signed rank, two-tailed). Statistical analyses were carried out in R (44).

## Supplementary information

Supplementary Table 1. Dataset of 1857 review articles.

## Acknowledgements

We would like to thank Britt Koskella, Craig MacLean, Kayla King, Divjot Kaur, Ravinash Krishna Kumar, Elisa Granato, Liam Shaw, Emma Burnett, and Michelle Pfeffer for their very helpful comments and discussions on this project. Certain data included herein are derived from Clarivate *Web of Science*. © Copyright Clarivate 2022. All rights reserved.

